# Single Cell RNA sequencing reveals transitional states and signaling shifts in nephron progenitor cells of the late-gestation rhesus macaque kidney

**DOI:** 10.1101/2025.08.18.670897

**Authors:** Kairavee Thakkar, Sunitha Yarlagadda, Lyan Alkhudairy, Andrew Potter, Konrad Thorner, Praneet Chaturvedi, Cristina Cebrian, Kyle W. McCracken, Nathan Salomonis, Raphael Kopan, Meredith P Schuh

**Author notes:** **Correspondence:** Meredith P Schuh, MD,3333 Burnet Ave, MLC 7022 Cincinnati, OH 45229.

## Abstract

**Background:** Human nephrogenesis is complete at 34-36 weeks gestation, with 60% of nephrons forming during the third trimester through lateral branch nephrogenesis (LBN). Currently, no mechanism exists for LBN as there are no late gestation human kidney transcriptional datasets. We hypothesized that a differentiated but dividing population of nephron progenitor cells (NPCs) would contribute to the amplification of nephrons in late gestation. We used the rhesus macaque, an established model of LBN, to help identify potential mechanisms.

**Methods:** Single-cell RNA-sequencing (scRNA-Seq) was performed on cortically-enriched fetal rhesus kidneys (n=9) from late second trimester and third trimester during LBN. This data was integrated with publicly available human scRNA-seq datasets from 8-18 weeks gestation kidneys (n=8) using state-of-the-art bioinformatics pipelines. Differentially expressed genes and ligand-receptor interactions were assessed and validated using RNAScope^TM^ on human and rhesus archival tissue.

**Results:** scRNA-Seq of 64,782 rhesus cells revealed 37 transcriptionally distinct cell populations, including 7,879 rhesus NPCs. Pseudotime analyses identified a late gestation-specific lineage branch of differentiated NPC in rhesus that was not observed in mid-gestation humans. Differential expression analyses identified increased *SFRP1, FZD4*, and *TLE2* and decreased *FZD7*, *SHISA2*, *SHISA3*, and *TLE4* within the late-gestation rhesus NPC compared to mid-gestation human NPC and increased SEMA3D within the rhesus ureteric bud (UB) tip, suggesting a compositional shift in WNT and SEMA signaling components within the naive NPC population during LBN.

**Conclusion:** The rhesus macaque uniquely enables molecular studies of late-gestation primate nephrogenesis. Our study suggests the hypothesis that a transitional state of self-renewing NPCs supported by compositional shifts in key pathways may underlie the switch from branching phase nephrogenesis to lateral branch nephrogenesis and support ongoing nephron formation in late gestation.

**Translational statement:** No transcriptional data exists for the late-gestation human kidney, during which 60% of the nephron endowment is formed through a primate-specific process called lateral branch nephrogenesis (LBN). In this study, we used single-cell RNA sequencing the late gestation rhesus macaque as a model for this developmental stage. Our findings suggest a potential mechanism for LBN, in which a transitional state of self-renewing nephron progenitor cells (NPCs) is supported by compositional shifts in key pathways, allowing for continued nephron formation in late gestation.

## INTRODUCTION

The period of new nephron formation, or nephrogenesis, in human kidney development typically extends through 34-36 weeks gestation (WG), with 60% of nephrons forming after 23 WG through a process termed lateral branch nephrogenesis (LBN).[1] Infants born prematurely must undergo LBN postnatally and are at the low end of nephron endowment and at high risk for chronic kidney disease (CKD) later in life.[2–10].In the mouse, nephrons are formed in a process associated with bifurcating branching of the ureteric bud (UB), which we will refer to as branching phase nephrogenesis (BpN) (**Fig 1A**). [11, 12] In contrast, human nephrogenesis transitions through three stages, from BpN (5-15 WG) to the short arcading phase (15-22 WG) **(Fig 1B)** to LBN (23-36 WG), where an elongating, non-branching UB tip induces nephrons, each directly connecting to the ureteric stalk after induction **(Fig 1C)**. While mouse models have been instrumental in elucidating the fundamental paradigms that control nephron progenitor cell (NPC) behavior and nephron formation, which appear to be highly conserved in the earliest stages of human kidney organogenesis,[13] they cannot provide insight into this critical LBN period of development that produces the majority of nephrons in humans and is aberrantly regulated in babies born preterm, affecting approximately 10% of all births or over 13 million infants born annually[14].

**Fig 1.**
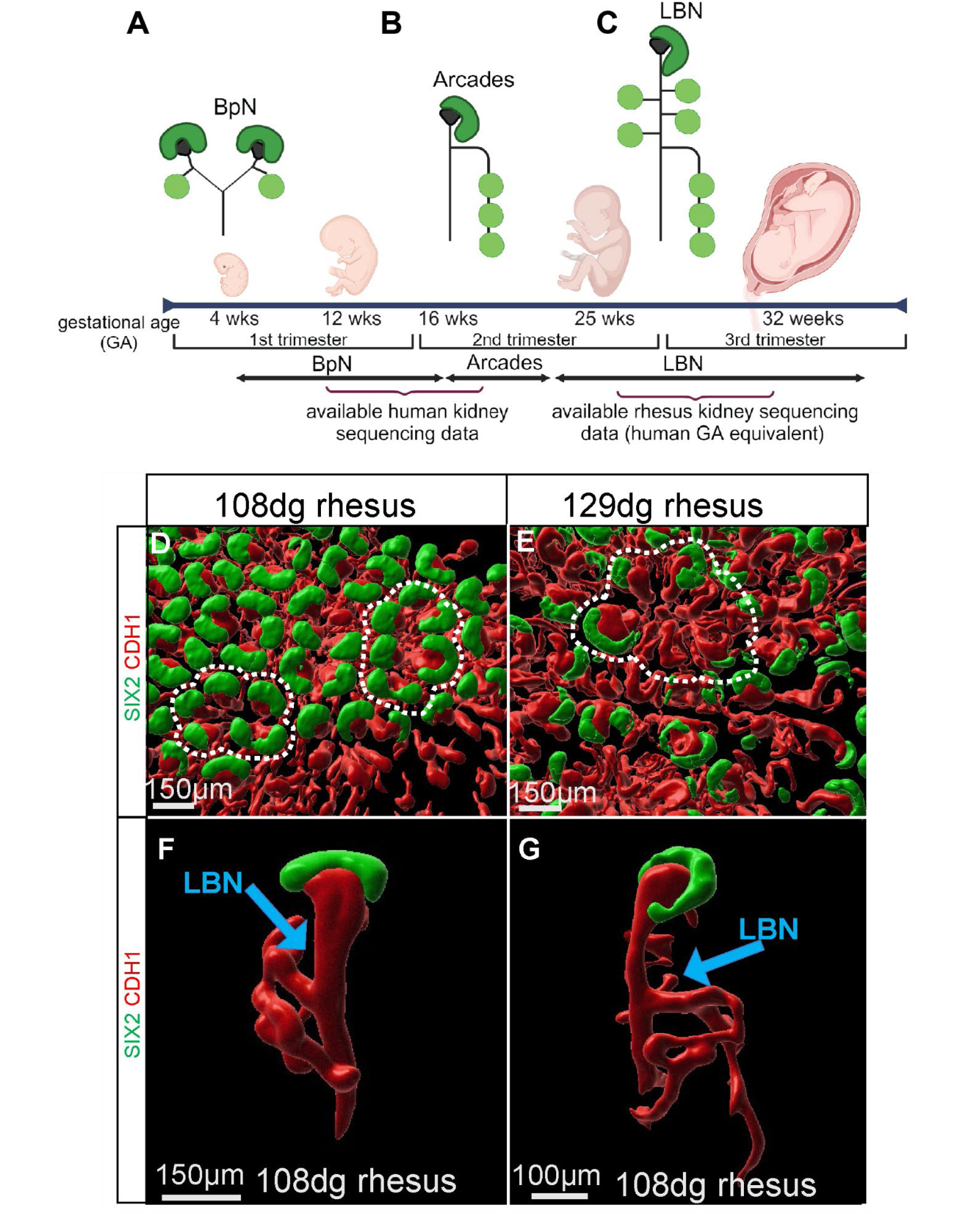
Stages of nephrogenesis in human and comparative rhesus morphology. **A)** Rhesus gestational period is 165 days and divided into three trimesters. During branch phase nephrogenesis (BpN), nephrons are formed in a process associated with bifurcating branching of the ureteric bud (UB). **B)** 15 weeks gestation (WG), human nephrogenesis transitions to arcading phase followed by **C)** LBN at 23-36 WG, where an elongating, non-branching UB tip induces nephrons. Rhesus gestation in days included for reference. **D)** LBN (108dg) rhesus displays the same rosette-like organization of the NPC niche as the **E**) 129dg rhesus. **F-G)** Further evidence of LBN during 108dg (late second trimester). Lateral branches (blue arrow) are seen along the ureteric stalk.

For many reasons, no transcriptional data exists for the late-gestation human kidney, and available human scRNA-seq datasets are limited to a developmental period before LBN begins. Additionally, human fetal archival material from autopsy can be challenging to obtain and can be subject to RNA degradation [15, 16]. To overcome these challenges, we established the rhesus macaque [17, 18] as an LBN model and identified age-dependent changes within the late gestation nephron progenitor cells[18] compared to early/mid-gestation human nephrogenesis **(Fig 1)**[19–21] and the fully differentiated kidney. Despite identifying changes within the nephrogenic niche, we were previously unable to identify signaling pathways in the original dataset due to the sparsity of NPCs nearing nephrogenesis cessation at ∼130dg (316 NPCs). Specifically, it remained unclear how self-renewing NPCs are maintained in a niche, comprised of a single non-branching UB tip, despite a known change in transcriptional signature[18] and an increase in WNT signaling over time[22].

In this study, we overcame the limitations of our previous investigation using integrative single-cell transcriptomics to perform a comprehensive and high-resolution comparison of nephrogenic niche progenitors during LBN and BpN, composed of 7,879 NPCs, to investigate how nephron amplification occurs independent of branching in the primate-specific phases of nephron production. With this comprehensive dataset, our findings support that a transitional state of self-renewing NPC, which exhibits features classically associated with an early induced state, contribute to ongoing nephron formation but also can also be maintained within the niche through changes in the signaling milieu. Given that late-gestation human and early-gestation rhesus samples are not accessible for ethical and logistical reasons, we performed cross-species analysis to bridge developmental stages of primate nephrogenesis from mid to late gestation. Our cross-temporal cross-species analysis identified that compositional changes in the WNT and SEMA3 signaling pathways between the NPC and UB may retain these cells that would otherwise differentiate at earlier stages of kidney development.

## METHODS

### Animals

All primate procedures were performed at the California National Primate Research Center at University of California Davis according to the Institutional Animal Care and Use Committee approved protocol #20330 awarded to Claire Chougnet. Additional information can be found in Supplemental Methods.

### Rhesus single-cell RNA sequencing

Rhesus genes were translated to human orthologs (using hg19, Ensmbl release 75). The intersection of genes observed between the human scRNA-seq datasets (see the next section in Methods) and the human orthologs observed in rhesus scRNA-seq datasets were considered for all downstream analyses.

### Human single-cell RNA sequencing

Count matrices of the human scRNA-seq datasets were downloaded from GSE114530, GSE112570, and GSE102596. We filtered the count matrices for only the cells that had a cell type annotation in the original publication. To keep cross-species comparisons and analyses consistent, we also filtered the count matrices to only those genes that were also seen in the human ortholog genes from the rhesus samples. These filtered count matrices from each sample were normalized and log-transformed using the *NormalizeData* function in Seurat and merged.

### Single-cell RNA sequencing analyses

#### Additional details can be found in the Supplemental Methods

#### Cell type annotation

SoupX adjusted counts from each rhesus sample were normalized and log-transformed using the *NormalizeData* function in Seurat and merged. To annotate cells in this dataset, we used the software cellHarmony to identify our previously published 37 transcriptional cell identities. The merged log-normalized file was provided as the input for cellHarmony. For the reference dataset, we used the scRNA-seq dataset and the 37 cell identities of rhesus kidneys available at GSE158304. Cell-identity pseudobulks of only the marker genes of the 37 clusters were calculated and provided as the reference for cellHarmony. The correlation cutoff for cell alignment was set to 0.8.

#### Subclustering NPCs and UB cells

Cells from clusters c25, c26, c28, and c29 were considered as NPCs and were further clustered using ICGS2, an unsupervised clustering algorithm. Default parameter values were set in ICGS2 (cosine clustering) in addition to exclusion of cell-cycle effects. Marker genes were identified for each NPC subcluster using the MarkerFinder algorithm (Supplemental Table 7). Cells within these NPCs that expressed any two out of *SIX1, SIX2* and *ITGA8* were considered as early transitional NPCs.

Cells from clusters c11 and c13 were considered as UB cells. UB stalk and tip clusters were identified in our previous publication (GSE158304) and we use these clusters and dataset as the reference. Then, we identified the 6 UB subclusters in the rhesus samples using cellHarmony. Cell-identity pseudobulks of only the marker genes of the six subclusters from the reference dataset were calculated and provided as the reference for cellHarmony. The correlation cutoff for cell alignment was set to 0.8.

Similar to the rhesus dataset, we applied cellHarmony on the human scRNA-seq data to identify the 37 transcriptional cell identities. For cellHarmony analysis, the reference file is the same as described for the rhesus cell type annotation, and merged log-normalized counts of human samples were provided as the input. Correlation cutoff was set to 0.8. Similarly, cellHarmony was used to project UB subclusters and NPC subclusters in human with relevant reference (pseudobulks of the cluster marker genes using using the log-transformed gene expression) from rhesus dataset.

Integrated UMAP of rhesus and human data was created using Harmony. Anndata objects containing the log-normalized gene expression of rhesus and human cells labeled by cellHarmony were merged. Mitochondrial genes were removed for downstream analysis. Using the merged Anndata object, top 3000 highly variable genes were determined using the Seurat flavor. Sources of variation due to total RNA counts and percentage of mitochondrial counts were regressed out. PCA for 50 components was performed using the the highly variable genes. Then, harmony analysis was done on this PCA using the harmony function from harmonypy package. Harmony-corrected PCA matrix was then used for generating UMAP using scanpy.

### Cross-species analyses of the early transitional NPCs

#### Differential gene expression analysis

Differentially expressed genes of NPCs and UB C6 in rhesus as compared to those in the human dataset were determined using the software cellHarmony, which calls a moderated t-test. Log-normalized gene expression matrices from human and rhesus were provided as the input. Genes with fold change greater than 1.2 and FDR-corrected p-value less than 0.05 were considered differentially expressed.

#### Lineage trajectory inference, pseudotime analysis, and Cell-cell signaling interactions

Pseudotime analysis was done separately on the rhesus and human datasets using Monocle2.

Cell-cell signaling interactions were determined individually for the human and rhesus datasets using the R package CellChat (v2.1.2). Additional information can be found within the supplemental methods.

### Human Kidney Validation Studies

Human kidney formalin fixed tissue samples were obtained from the Cincinnati Children’s Hospital Medical Center Biobank after review and approval by the Internal Review Board (IRB) for use of Discover Together biobank. Further details can be found in the supplemental methods. Due to poor quality of biobank mid-gestation samples, we were unable to perform RNAScope^TM^ at mid-gestation timepoints, and only late-gestation human samples were used for RNA in situ hybridization validation studies.

## RESULTS

### Lateral branch nephrogenesis in the human and rhesus is characterized by elongation of a single active ureteric tip that induces nephrons without branching

The gestational period of the rhesus macaque is 165 days and divided into three trimesters. We previously evaluated the morphology of third-trimester (∼130 days gestation (dg)) rhesus kidney, which is equivalent to 32 weeks gestation in the human.[18]. In this study, we obtained rhesus fetal kidney tissue from two donors at 104-108dg (∼25-26 WG in the human or late second trimester) (**Fig 1D-G**).[18] Like the ∼130dg rhesus samples, the 105-108dg samples demonstrated rosette-like organization of the nephrogenic niche as well as LBN morphology (**Fig 1D, E**). LBN can be identified by the SIX2+ NPC cap associated with a single UB tip, with CDH1+ lateral branches extending from the ureteric stalk below (**Fig 1F,G blue arrow)** supporting the use of data from late second trimester as a rich source of NPC during LBN. These findings of LBN in late second trimester correspond appropriately to morphologic human data where LBN begins at 22WG[23]. A schematic of how nephron induction may occur during LBN in late second trimester samples in rhesus and human in the absence of UB branching is shown in **Supplemental Figure 1**.

### Single-cell RNA-sequencing reveals late-gestation specific progenitor cell populations undergoing lateral branch nephrogenesis

We hypothesized that comprehensive molecular characterization of the niche progenitors through transcriptomic studies would uncover the mechanisms underlying this distinct behavior of nephrogenesis uncoupled by UB branching. We supplemented our scRNA-seq dataset of four fetal rhesus kidneys at ∼130dg[18] with five additional rhesus samples, including two samples at 105-108dg (∼7 times more progenitor cells per sample), for a total of nine cortically-enriched fetal kidney samples from distinct donors (**Supplemental Table 1**). We identified 37 transcriptionally distinct cell clusters from 64,782 cells using our previous cluster annotations as reference (**Supplemental Fig 2A**, **Supplemental Table 2, 3**).[18] Due to lack of available data from mid-gestation rhesus or late-gestation human samples, we integrated our scRNA-seq data from late-gestation rhesus with that available from human fetal kidneys (GSE114530[19], GSE112570[20], and GSE102596[21]) from seven samples (representing BpN and arcading), consisting of 29,484 cells, using Harmony (**Fig 2A, Supplemental Fig 2B, Supplemental Table 4**).[24] Four clusters (c25, c26, c28, c29) were marked with definitive NPC markers such as *SIX1*, *SIX2*, *CITED1* and *MEOX1*, with cluster 25 containing the naive NPCs (in rhesus, n = 7879 cells; in human, n=9434 cells) (**Fig 2B**). UB markers identifying clusters 11 and 13 included *WNT11*, *WNT9B*, *GATA3*, *ADAMTS18*, and *ELF5* (**Supplemental Fig 2C**). To specifically interrogate and esolve the niche progenitor states, we subclustered the four NPC and two UB progenitor clusters using ICGS2, an unsupervised non-negative matrix factorization-based clustering pipeline, which yielded 8 subclusters of NPCs and 6 subclusters of UB progenitors (**Fig 2C**, **Supplemental Table 5, 6**). NPC C4 represented the most naive NPC subcluster, with high expression of *SIX1, SIX2, MEOX1, EYA1, CITED1, ITGA8, and TMEM100* (**Fig 2D, Supplemental Table 7**), while the other NPC clusters represent different states of proliferation and commitment to differentiation.

**Fig 2.**
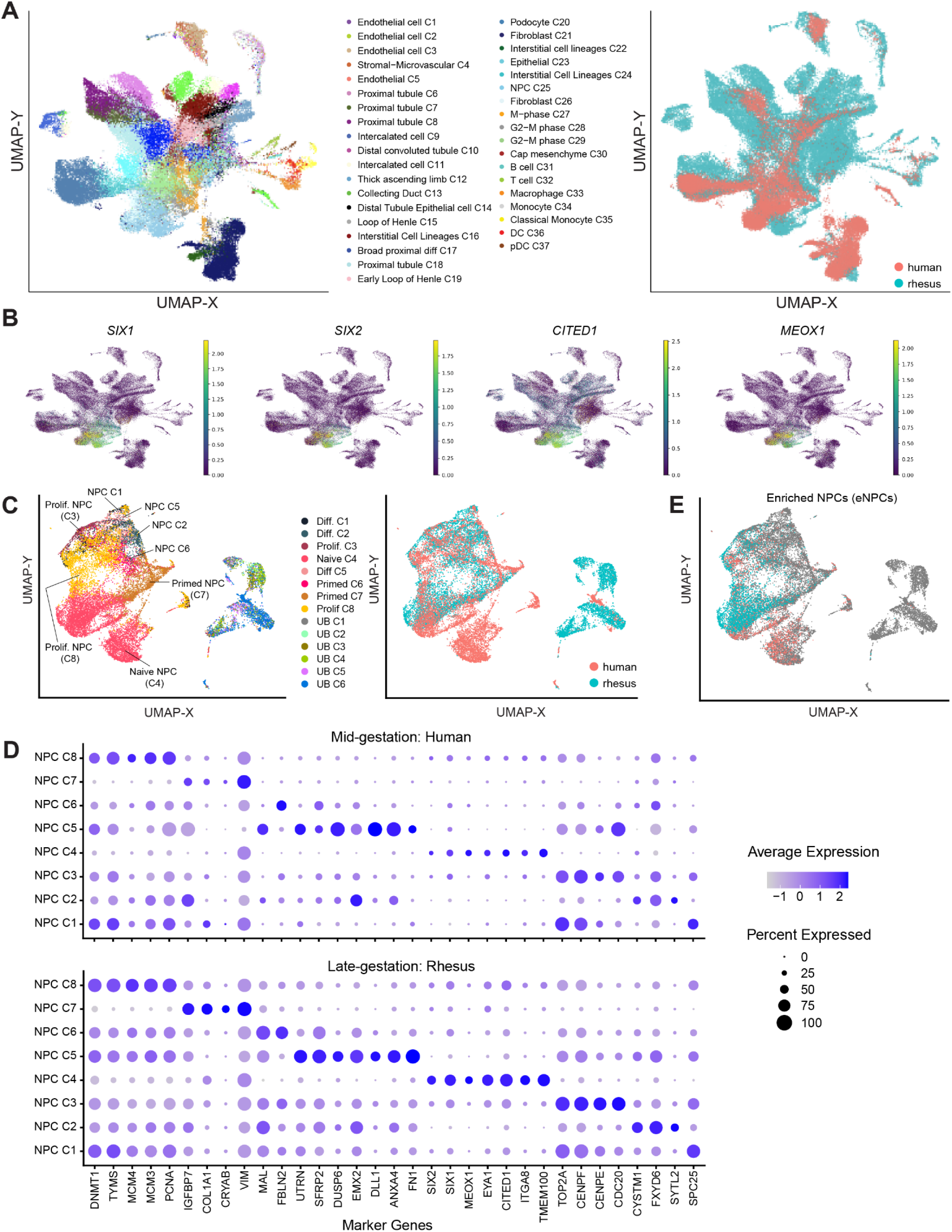
Cross-species integrative single-cell transcriptomics characterizes heterogeneity and temporal differences in NPCs. **A)** Integrated UMAP of 94,266 cells from cortically-enriched fetal kidneys from 9 rhesus (late second and third trimesters) samples and 5 human (8-13 weeks old) samples. Cells are colored by cellHarmony cluster labels (left panel) and by species (right panel). **B)** UMAP from panel A colored by gene expression of key NPC progenitor marker genes. **C)** Integrated UMAP of NPC and UB cells from rhesus and human samples. Cells are colored by cell subclusters within NPCs and UB (left panel) and by species (right panel). **D)** Dot plot indicating average expression of major marker genes of NPC subclusters in human (top panel) and rhesus (bottom panel) datasets. Dots are sized by the percent of cells that expressed the marker gene. **E)** UMAP from Panel C highlighting the enriched NPCs (eNPCs) defined by expression for: *SIX1*, *SIX2*, *ITGA8* (at least 2 of the three genes expressed). The highlighted cells are colored by species, similar to right panel C.

To reduce the likelihood that comparative analyses between rhesus and human would be confounded by committed NPC, we filtered the NPCs to include those that expressed at least two of the NPC markers: *SIX1*, *SIX2*, and *ITGA8 (*referred as enriched NPC, or eNPC, hereon)(**Fig 2E**). In rhesus, 80% of these 2,719 eNPC came from the 105-108dg samples (**Supplemental Fig 3, Supplemental Table 5, 6**), further supporting the utility of these newer samples from our previous investigation. *CITED1* was not chosen as a marker for filtering due to its relatively broad expression throughout the rhesus cells (**Fig 2B**). After filtering for eNPC, 64.5% (previously 31.5%) and 93.9% (previously 55.8%) of the cells were from NPC C4 in rhesus and human, respectively (**Supplemental Fig 3**). To investigate the characteristics of the removed human C4 cells, we performed differential gene expression of eNPCs compared to the removed cells. We noted increased expression of *SIX1*, *SIX2,* and *ITGA8* in the eNPCs, supporting this subset as the most naïve progenitors while maintaining a sufficient number of cells in C4. Cells removed demonstrated upregulation in genes related to heat shock responses (HSPA1A, HSPA1B, HSPB1, HSP90AA1, TCIM), potentially representing differences in dissociation protocols. (**Supplemental Table 7**). Interestingly, we noted three subclusters (C1, C5, C6) in the rhesus that remained after filtering (**Supplemental Fig 3A**), supporting their identification as late-gestation specific progenitors (**Fig 2D, Supplemental Table 7**).

### Late-gestation NPC show a distinct trajectory from mid-gestation human NPC

To orthogonally confirm that C4 cluster represented the most naive NPC state and to delineate the differentiation trajectories using an unsupervised approach, we used Monocle[25] to perform pseudotime analysis of the eNPCs at mid gestation (human) and late gestation (rhesus) (**Fig 3**). Branch points along the constructed cell trajectory mark a suggested split into two or more distinct cell fates. In the mid-gestation human (**Fig 3A**), the first and only branch (which we will refer to the conserved primate branch) splits the NPC C4 into more differentiated trajectory (left) or a proliferative state (right) (**Supplemental Table 7**). Despite having a comparable number of C4 NPCs in rhesus and human, the pseudotime analysis identified an additional branch (late-gestation rhesus branch) in late-gestation rhesus NPCs not seen in the early human samples (**Fig 3B)**. This initial (late-gestation rhesus branch creates two NPC populations: a transitional but terminal right sided branch consisting of naïve and differentiated NPC (NPC C4 and C8) and a left branch that later gives rise to the conserved primate branch (C4).

**Fig 3.**
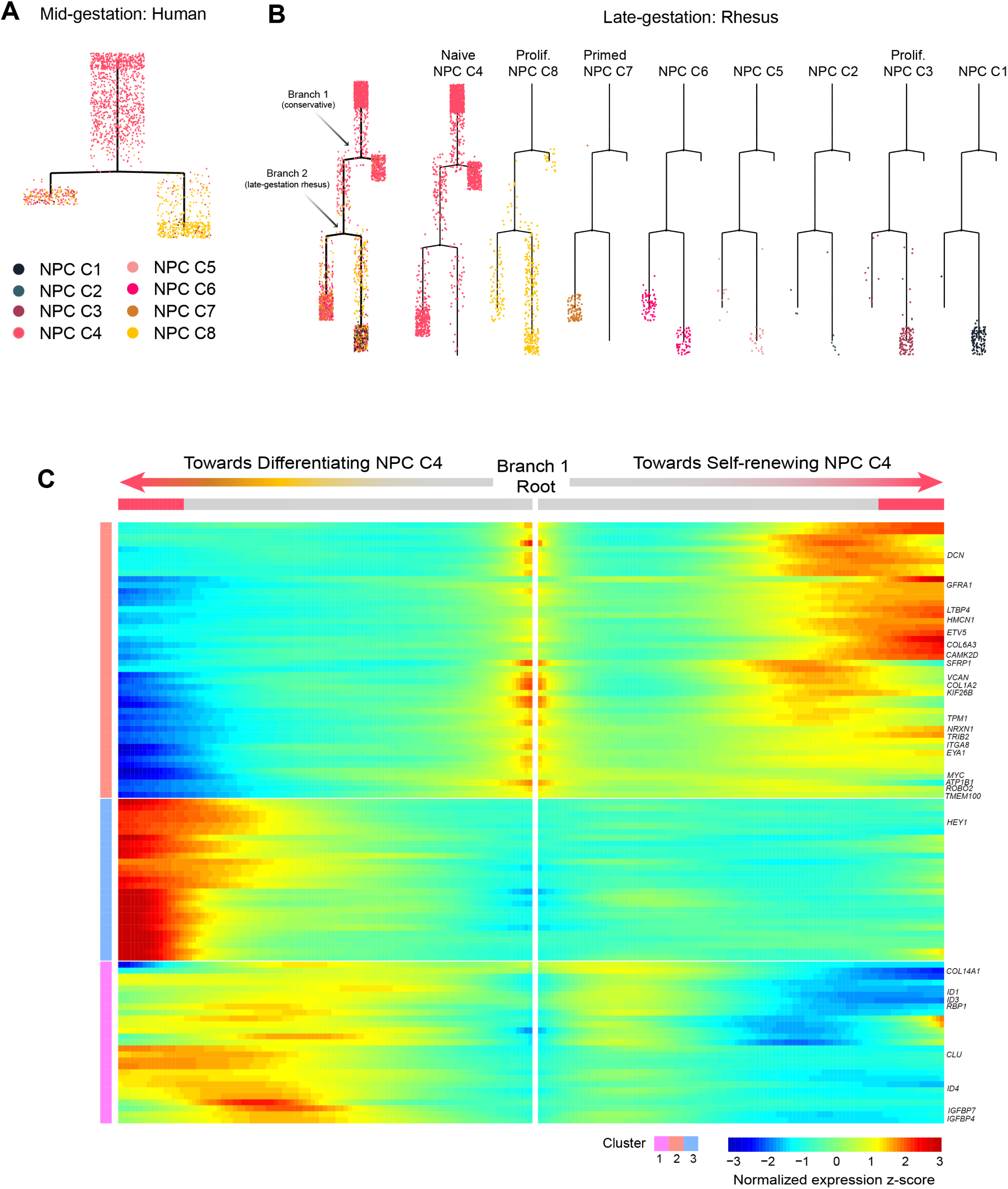
Developmental trajectory of NPCs. Lineage trajectory of NPCs constructed by Monocle2 for **A)** human and **B)** rhesus. Pseudotime increases from top to the bottom of the tree. To clearly illustrate where clusters are located, the lineage tree is reproduced to highlight each rhesus cell population. **C)** Heatmap of normalized gene expression showing branch-dependent genes (Monocle BEAM test; FDR < 10%) as rows along the two fates of Branch 1 in rhesus. Left and right segments show the gene expression change along the left and right branches, respectively, after Branch 1 in the Monocle tree in panel B. Columns are points in pseudotime, and the dividing middle segment of the heatmap indicates beginning of pseudotime. Selected genes are highlighted on the right. Rows are clustered using hierarchical clustering.

To further understand this unexpected lineage bifurcation, we examined the transcriptional differences in the rhesus C4 cells using Monocle BEAM analysis and published literature.[20, 26, 27] (**Fig 3C, Supplemental Fig 4**) The rhesus-only branch distinguishes the differentiating eNPCs (left) vs self-renewing eNPCs (right). This right-sided trajectory includes NPC markers *EYA1*, *TMEM100, NRXN1, ITGA8*, and *RSPO3*, as well as what has previously been referred to as primed NPC[20], including *SFRP1*, *ROBO2*, and *TPM1*. However, this branch also contains markers of proliferation (*ATP1B1, CAMK2D*), proximal (*MYC*), medial-distal (*TRIB2*), and distal segment (*LTBP4, PCDH18, COL6A3*). Additionally, these progenitors include markers typically associated with extracellular matrix/interstitial progenitors including *COL1A2*, *DCN*, *VCAN*, and basement membrane markers *COL14A1* and *HMCN1*. (**Fig 3C, Supplemental Table 8**). In the left (more differentiated) trajectory of the rhesus-only branch, genes included primed NPC markers *HEY1, ID1, ID3*, and *ID4*, proliferation markers *TOPS2A, NUSAP1, MKI67, CDC20*, and early renal vesicle/broadly proximal markers *JAG1, HES4, SFRP2* and distal markers *CCND1* and *BMP7*. Interestingly, both sides expressed early proximal marker CDH6 and podocyte marker *ADAMTS9*. These data indicate that the left-sided population would be the one giving rise to more differentiated NPC populations and derivatives via established developmental paradigms. These findings suggest a transitional state of NPC in late-gestation that does not exist in mid-gestation and may provide the source of self-renewing NPC in late-gestation despite their induced state.

### Enriched NPCs present age-dependent differences in *WNT* signaling

To evaluate the age-dependent changes in NPCs, we performed differential expression of the NPC subclusters in rhesus vs human (**Fig 4A, Supplemental Table 9**). Genes prominent in the WNT pathway were identified within the NPC, including Frizzled (FZD) receptors, WNT inhibitors, and Transducin-Like Enhancer of Split (TLE) co-repressors (**Fig 4B)**. Specifically, we identified increased *FZD4* and *TLE2* (C7) and decreased *FZD7 (C3), SHISA2 (C4, C7), SHISA3 (C4), and TLE4 (C4, C7, C8)* **(Fig 4A-B**). We noted an increase in secreted Frizzled-related protein-1 (*SFRP1)* in the late-gestation rhesus relative to mid-gestation human, strongest within naïve NPC (C4) but present throughout the late-gestation NPC population, consistent with our findings of increased *SFRP1* within the self-renewing NPC branches of the Monocle pseudotime analyses. We also noted increased *SFRP2* in the more differentiated NPC clusters C2,3,5, and 6.

**Fig 4.**
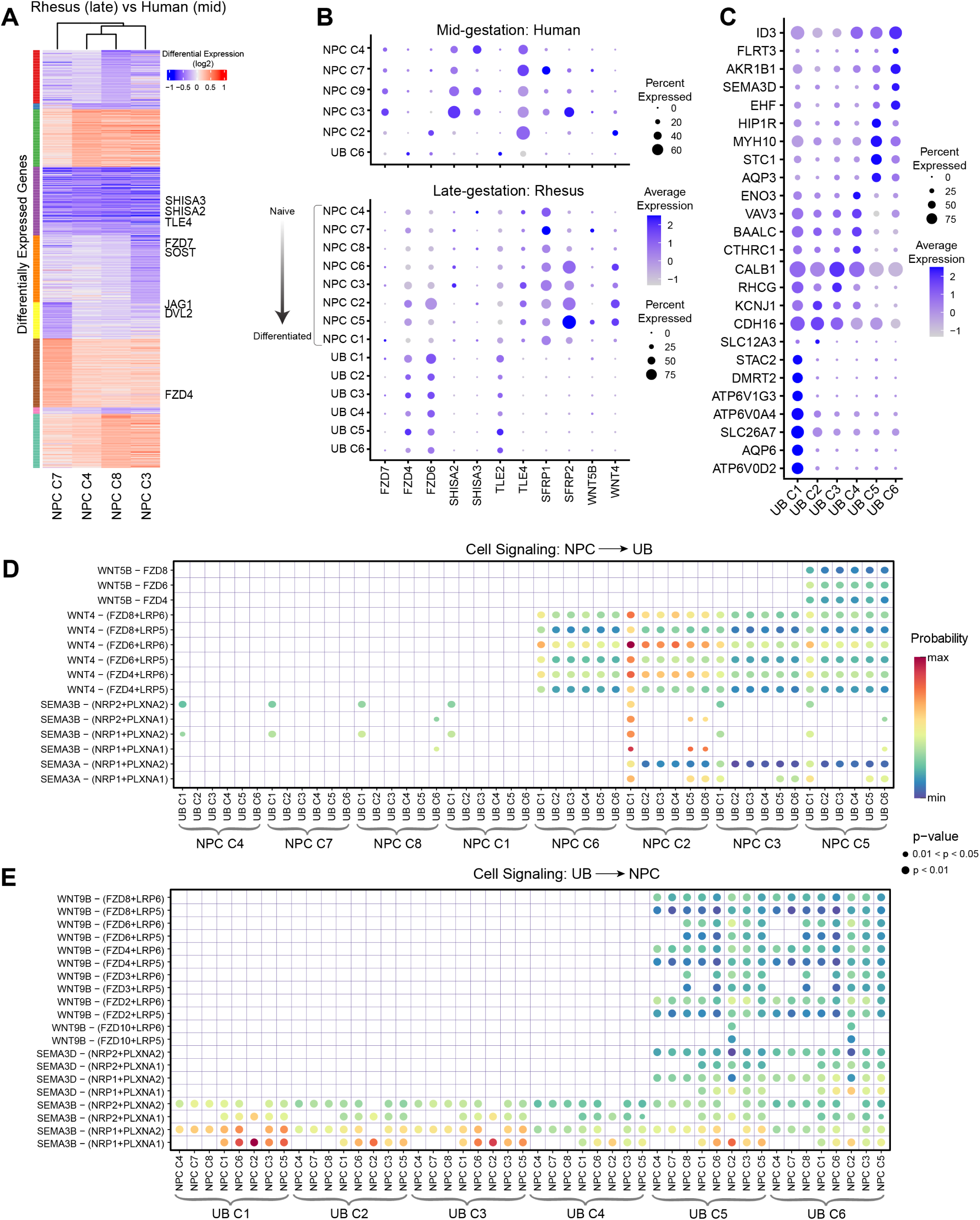
Age-dependent changes in WNT signaling. **A)** Heatmap indicating the log2 fold changes of differentially expressed genes (FDR p-value < 0.001) of each NPC subcluster in rhesus as compared to human cells. Top 150 differentially expressed genes are considered from each comparison. **B, C)** Dot plot indicating average expression of **B)** selected genes associated with WNT signaling in NPC subclusters in human (top panel) and rhesus (bottom panel) datasets, and **C)** marker genes of UB subclusters in rhesus. Dots are sized by the percent of cells that expressed the gene. **D, E)** Bubble plots showing the WNT-associated ligand-receptor pairs (rows) that contribute to secreted signaling **D)** from NPC to UB and **E)** UB to NPC in rhesus. Dots are colored by communication probability and sized by p-value.

The mechanisms controlling nephrogenesis involve cross-talk of signaling pathways between the NPCs and the closely associated UB tips (**Supplemental Fig 1**). Our initial subclustering (**Fig. 2C**) of UB cells (clusters c11 and c13) from the integrated dataset identified four stalk clusters (C1, C2, C3, C4) and two tip clusters (C5, C6), consisting of 3,507 total cells in rhesus and 391 cells[19] in human (**Fig 2C, Supplemental Table 10**). Over 95% of the UB cells from the human dataset were classified as UB C6, which limited the temporal comparative analyses to only UB C6 among the UB cells. Supporting a possible reduction in WNT response in the UB tip, we identified overall low expression of AXIN2 in the rhesus UB clusters (**Supplemental Figure 5A, Supplemental Table 11)**.

To further understand the signaling between eNPCs and UB, we interrogated the possible secretory receptor/ligand interactions between eNPC and UB in mid vs late gestation using CellChat.[28] Within the late-gestation rhesus, we identified distinct compositional changes within the WNT pathway (**Fig 4D-E, Supplemental Figure 5B, Supplemental Table 12**). While both human and rhesus exhibited WNT4 signaling to FZD4/6/8 from the more differentiated eNPC (C2), the rhesus NPC subcluster C5 demonstrated WNT signaling to UB tip and stalk through WNT5B ligand and FZD 4/6/8 interaction (**Fig 4D, E**).). Combined with the overall higher expression of the secreted antagonist SFRP1 in the naïve NPCs of rhesus, these data suggest an important role for differential modulation of WNT signaling activity as a mechanism governing LBN morphogenesis, consistent with the known role of the pathway as a pivotal regulator of kidney organogenesis.[22, 29–33] To further validate these findings, *in situ* hybridization was used to spatially localize the expression of differentially expressed components within the niche. We identified co-staining of *SFRP1* with *SIX1* in the cap NPC population (**Fig 5A**) that extends to the pretubular aggregates (PTA) and renal vesicles (RV) in late-gestation rhesus and human. This staining pattern confirmed our DEG analysis within the late-gestation rhesus and supports our pseudotime gene expression changes in the C4 rhesus progenitor population. *SFRP1* extended also into the surrounding stroma in addition to the NPC. *SFRP2* was expressed in the early *SIX1+* differentiated nephron structures (PTA/RV), consistent with its increased transcriptional level in the induced NPC *WNT4*+ population of cells.[22, 34]. We quantified these differences within the nephrogenic niche and identified more *SFRP1* compared to *SFRP2* in the NPC, supporting the above findings (**Fig 5B**). This relationship was maintained when normalizing to *SIX1* expression.

**Fig 5:**
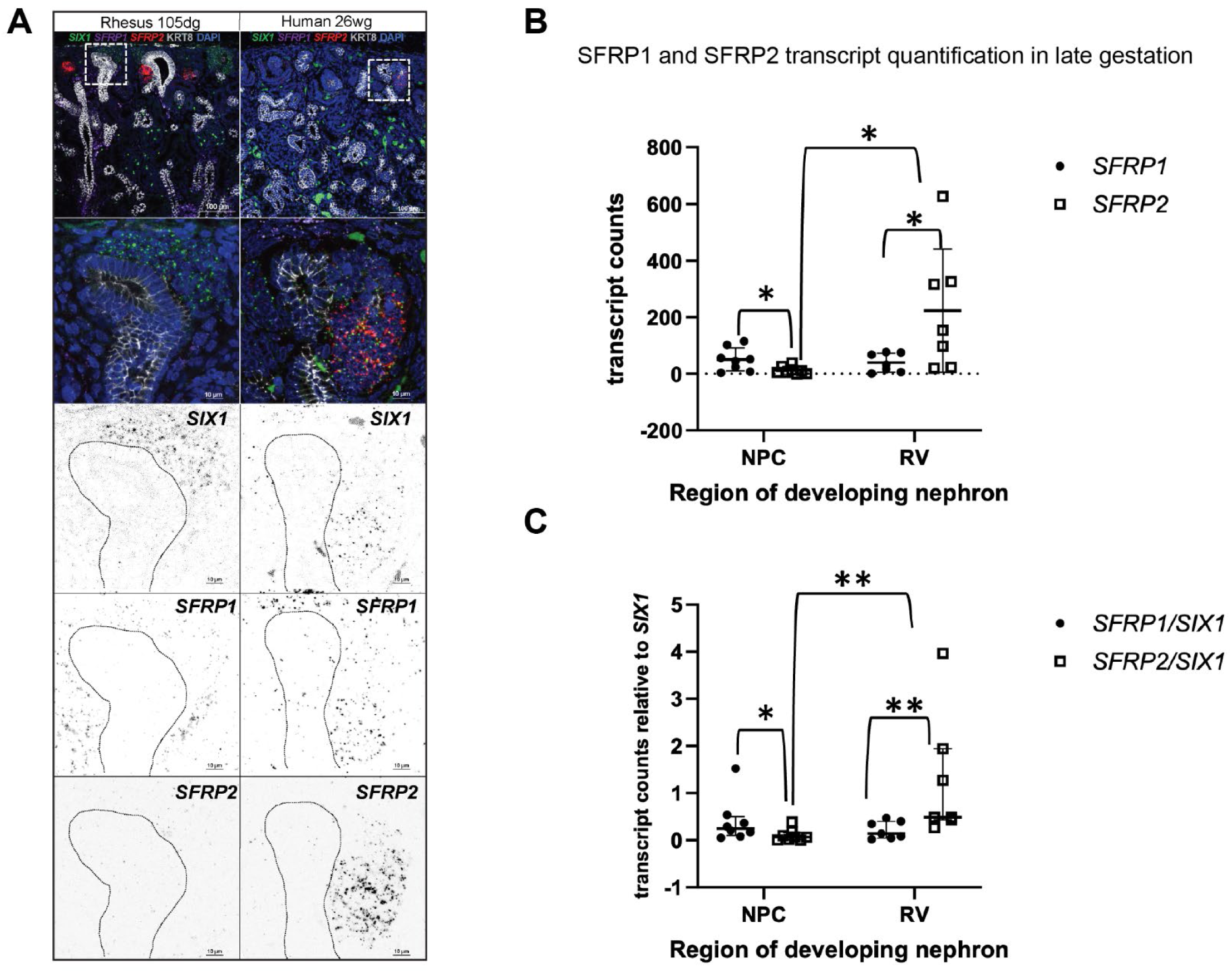
RNAScope^TM^ validation of *SFRP1* expression in late gestation. **A)** Representative late-gestation rhesus and human kidney with co-localization of SFRP1 (purple) and SIX1 (green), with additional SFRP1 expression within the surrounding stroma. SFRP2 (red) was noted further away from the cortical surface. **B)** Quantification of SFRP1 and SFRP2 total transcripts in nephron progenitor cells (NPC) and renal vesicle (RV) as absolute counts. Mean and standard deviation presented on plot. **C)** SFRP1 and SFRP2 relative to SIX1. Data was not normally distributed, so median and interquartile range presented on plot.

We next also noted a change in *FZD* composition within our DEG and highlighted in the cell chat analysis. Based on DEG, we anticipated high expression of *FZD4* and low expression of *FZD7* in late gestation compared to mid gestation. On tissue validation, we identified diffuse staining of FZD4 in both the UB and NPC (**Fig 6A**). *FZD7*, while still present in the NPC, was most prominent in connecting segment of the distal nephron fusing to the UB tip (**Fig 6A**), which was not anticipated based on DEG. This unexpected localization was also seen at the protein level in the human archival sample, with increasing FZD7 protein expression at the distal fusion points with increasing gestational age, and present in examples for arcading (17wg) and LBN (27-30wg) (**Fig 6B**). Returning to the larger dataset, we identified increased *FZD7* in cluster 15 in late gestation. Although this cluster is labeled as Loop of Henle (with marker genes including *MAP1B*, *TFAP2B*, and *IRX2*), there is some overlap of these genes with collecting duct and urothelial expressions, including *FRAS1*, *TSPAN13*, *LLGL2*, *CPEP2* and *PTPN3*, suggesting broad expression of *FZD7* from Loop of Henle to collecting duct epithelium during the fusion of the nephron to the ureteric stalk. Given that decreased *SHISA2* and *SHISA3* were identified on our differential expression analysis we next evaluated the expression of these two genes **(Supplemental Fig 5C)**. Despite the low expression in the DEG analysis, we identified strong *SHISA2* within the NPC and early differentiating structures at late gestation. Interestingly, *SHISA3*, while still present in the NPC, appeared most prominent in the surrounding stroma (**Supplemental Fig 5C**). *TLE2* and *TLE4* were also evaluated using RNAScope^TM^. *TLE4* was present within the NPC of late-gestation rhesus and human, while no distinct pattern was visible in *TLE2* expression (**Supplemental Fig 5D**).

**Fig 6.**
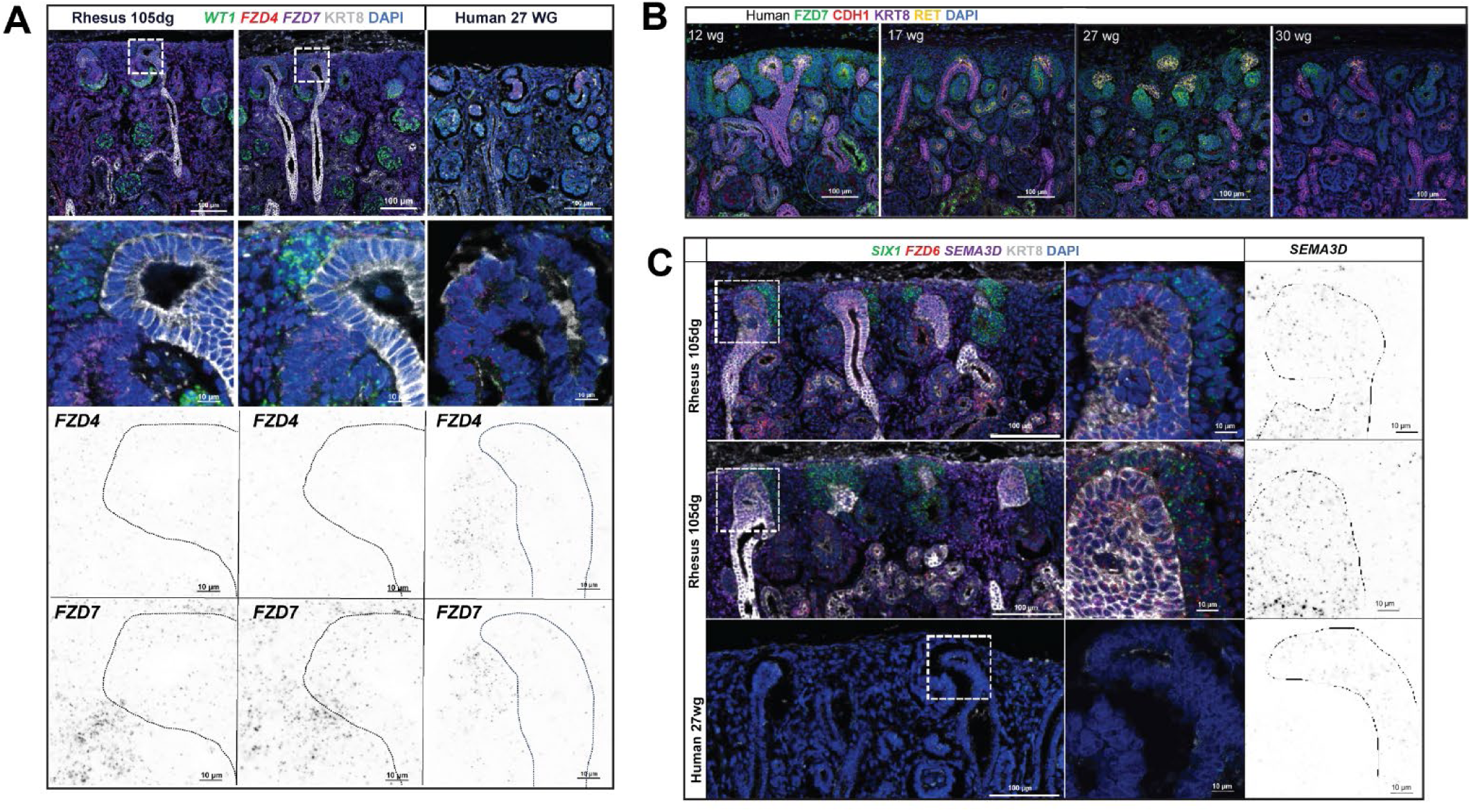
RNAScope^TM^ validation of WNT pathway and SEMA3D. **A)** FZD4 and FZD7 expression in the late-gestation rhesus and human. While FZD4 was broadly expressed in the NPC and developing nephron, FZD7 localized to the connecting segment or the distal nephron and fusion to the UB in rhesus (two niches shown) and human. **B)** FZD7 protein expression in the developing human kidney. FZD7 is seen at the fusion of the nephron segments in examples of arcading (17 wg) and lateral branch nephrogenesis (27-30 wg). **C)** Late-gestation rhesus and human demonstrate strong *SEMA3D* signal in the outer UB and UB tip, with additional SEMA3D staining identified within the maturing interstitium.

### Enriched UB cells present age-dependent differences in *SEMA3* signaling

SEMA3D expression was significantly upregulated in UB C6 of late-gestation rhesus compared to mid-gestation human samples (FDR p-value < 0.05) (**Supplemental Table 11**). Using CellChat, we identified SEMA3A and SEMA3B signaling from the differentiated eNPC via receptors NRP1/2:PLXN1/2 in the late gestation rhesus samples. We next interrogated the possible secretory receptor/ligand interactions from the UB to eNPC in the late gestation rhesus which suggested reciprocal semaphorin signaling in the niche (**Fig 4E**). SEMA3B ligand signaling to the NRP1/2 and PLXN1/2 receptors was observed ubiquitously across eNPCs.. In contrast, SEMA3D signaling to these receptors originated specifically from the UB tip clusters. We validated this UB tip marker marker using RNAScope^TM^.(**Fig 6C)**.

## DISCUSSION

In summary, the gene expression approaches performed in this study help overcome current limitations in identifying pathways acting late in development, as the direct study of late-gestation human kidney development is not possible. The rhesus is the only resource that can be leveraged to define critical cellular and signaling changes that occur during human kidney development at single-cell resolution. Through our cross-time, cross-species investigation of the late-gestation rhesus macaque compared to mid-gestation human, we identified molecular differences in the nephron progenitor cells, including distinct NPC clusters in late gestation and distinct trajectories of the naïve NPC in late gestation. Despite known limitations of the available samples, our data suggests a transitional NPC state supported by a potential change in the signaling milieu between the NPC and UB within WNT and SEMA3 signaling pathways, allowing for this differentiated but self-renewing NPC to be maintained within the niche. While we acknowledge the limitations of RNA analyses, this hypothesis provides a potential mechanism for nephron amplification during LBN.

### Distinct progenitor trajectories are seen within late-gestation rhesus

Distinct trajectory of the most naïve NPC in late gestation suggests there may be a population of late-gestation that does not exist mid-gestation NPC; a population of early differentiated NPC that contribute to self-renewal and remain in the niche. Rather than a single lineage branch dividing the self-renewing NPC from the differentiating NPC, the rhesus instead has two lineage branches. This right trajectory maintained strong cell cycle signals, markers of the naive NPC, as well as markers associated with the interstitial progenitors and early basement membrane markers, suggesting a shift in both location and early function despite their naive state that may be needed for LBN-nephrogenesis. This transitional proliferating population is distinct from those cells that go on to fully differentiate and become nephrons. The cells undergoing LBN may remain in a high cell cycle and early committed cell state (expressing early BpN markers). Slowing of cell proliferation has been associated in the aging of the progenitor niche[35, 36]. However, changes in cell cycle expression at the RNA level are not the full story, as low translation in stem cell populations, independent of cell cycle, have been shown to maintain the progenitor state compared to high translation in differentiated cells.[37] Given that these populations of cells still maintain their progenitor signature, they may represent the LBN and/or arcading phase of a self-renewing but early differentiated cell that can contribute to nephron formation in late gestation.

### Changes in WNT signaling in late gestation may drive the transition from BpN to LBN

One of the potential pathways identified through this investigation included alterations within WNT signaling in late gestation compared to mid gestation. Jarmas et al found that the “tipping point” for nephron progenitor exit from the niche is controlled by the gradual increase in the translation of WNT agonists in individual nephron progenitor cells, enhancing the response to ureteric bud-derived WNT9b inputs.[22] It remains unknown how this shift in WNT responsiveness with age could impact ongoing nephrogenesis in a model with prolonged LBN prior to nephrogenesis cessation. SFRP1 is a secreted WNT antagonist that interacts directly with WNT ligand and can block its interaction with receptors.[38–41] Our data suggests that increased expression of SFRP1 within the naïve NPC may shield these nephron progenitor cells from the action of WNT agonists, as suggested by low expression of downstream target AXIN2, allowing for prolonged, ongoing nephrogenesis in late gestation without branching. Without early gestation rhesus samples for a within-species analyses this remains speculative. Future experiments including overexpression of *SFRP1* in a genetically tractable model of the NPC population may shed light on this question.

*FZD7* identified the connecting segment/fusion of the distal nephron to the UB rather than the NPC population itself in late gestation. However, mid-gestation human data supports its expression throughout the NPC, as well as podocytes and loop of Henle, but only faintly expressed in the connecting tubule.[42] Our single cell data also supports expression within the Loop of Henle, despite tissue validated presence at the fusion of the developing nephron. FZD7 has also been identified as putative receptor for WNT6, which leads to de novo tubulogenesis in kidney epithelial cells in culture,[43] but its role within the fusion of the nephron is not known. FZD7 can interact with LRP5/6 to transmit canonical Wnt signaling, as well as through ROR2 to transmit non-canonical Wnt signals.[44] WNT5A, with potential signaling highlighted within the rhesus-only clusters, may play a role in this distal fusion during LBN. WNT5A non-canonical signaling through ROR2 plays an important role in mediating cell proliferation and migration within the metanephric mesenchyme around the wolffian duct[45], further supporting this hypothesis.

### The role of semaphorin 3D in UB elongation and induction of lateral branch nephrogenesis

Semaphorins are guidance proteins, mediating short and long-range repulsive and attractive guidance that influence organogenesis and disease.[46] Explant kidney cultures exposed to Sema3a demonstrate reduced ureteric branching, reduced GDNF expression, and reduced Ret phosphorylation. Conversely, Sema3a knockdown/knockout models resulted in increased bud branching.[47] Our study supports that *SEMA3D* is present within UB tips, as well as the surrounding interstitium, while our Cell chat data supports that *SEMA3B* signaling is prominent in the stalk. Our data suggests that increased *SEMA3D* signaling within the UB tip supports elongation rather than bifurcating branching, but future studies comparing the balance of *SEMA3B* to *SEMA3D* in the stalk vs tip require further investigation in early to late gestation. If early to mid-gestation tissue in the rhesus were available in the future, we would perform similar trajectory analyses of the UB overtime to better capture these changes.

### Limitations and future directions

Our study highlights a potential transitional, proliferating NPC that amplifies nephrons in late gestation LBN, which is the critical first step to maintaining these pathways otherwise impacted by preterm birth. However, the concepts proposed in this study remain hypothetical. Several limitations exist, resulting in unanswered questions. One of the greatest questions of this analysis is whether the identified changes within the late gestation NPC are time-dependent or species-dependent. No transcriptional data exists for the late-gestation human kidney, and available human scRNA-seq datasets are limited to a developmental period before LBN begins. Additionally, human fetal archival material from autopsy can be challenging to obtain and can be subject to RNA degradation.[15, 16] Future work will seek to define signaling pathways in the rhesus at mid-gestation before LBN begins, with validation studies at both translational and protein levels to further test these hypotheses. Acquisition of these earlier gestation rhesus datasets will allow us to answer the many questions that remain when defining the unique maturational changes in late gestation and provide the high tissue samples for spatial validation of our computational findings.

## ACKNOWLEDGEMENTS

The authors thank Dr. Claire Chougnet, Dr. Alan Jobe, and the California National Primate Center for donating the rhesus kidneys used for this study. This research was made possible, in part, using the Bio-Imaging and Analysis Facility (RRID:SCR_022628). and the Single Cell Genomics Facility (Kelly Rangel, Shawn Smith, RRID: SCR_022655). This study used samples from the Discover Together Biobank at Cincinnati Children’s Research Foundation. We thank the Discover Together Biobank for supporting this study, as well as participants and their families, whose help and participation made this work possible. We thank Katrina Schlum for initial insights into the data.

## AUTHOR CONTRIBUTIONS

R Kopan and MP Schuh conceptualized and planned the experiments, K Thakkar, R Kopan, and MP Schuh analyzed data. S Yarlagadda, L Alkhudairy, A Potter, and MP Schuh performed the experiments including single-cell dissociation. K Thakkar, N Salomonis, P Chaturvedi, and K Thorner performed bioinformatics analysis in collaboration with MP Schuh. K Thakkar and MP Schuh assembled figures. K Thakkar, KW McCracken, C Cebrian, and MP Schuh wrote the manuscript. The final manuscript incorporated input from all authors.

## DATA AVAILABILITY

The raw and processed files of scRNA-seq data from rhesus samples are deposited in GEO (Accession number: GSE305708). scRNA-seq data from human samples are available at GSE114530, GSE112570, and GSE102596. Code and scripts for reproducibility are available at https://github.com/kairaveet/integrative-nephrogenesis.

